# Interactions between insect vectors and plant pathogens span the parasitism-mutualism continuum

**DOI:** 10.1101/2022.09.27.509661

**Authors:** Ma. Francesca M. Santiago, Kayla C. King, Georgia C. Drew

**Affiliations:** Department of Biology, University of Oxford, Oxford, UK

**Keywords:** vector-borne, phytopathogen, multi-trophic interactions, symbiosis continuum, host fitness, vector-pathogen interactions, parasitism-mutualism continuum

## Abstract

Plants infected with vector-borne pathogens can suffer severe negative consequences, but the extent to which phytopathogens affect the fitness of their vector hosts remains unclear. Evolutionary theory predicts that selection on vector-borne pathogens should favour low virulence or mutualistic phenotypes in the vector, traits facilitating effective transmission between plant hosts. Here, we use a multivariate meta-analytic approach on 115 effect sizes across 34 unique plant-vector-pathogen systems to quantify the overall effect of phytopathogens on vector host fitness. In support of theoretical models, we report that phytopathogens overall have a neutral fitness effect on vector hosts. However, the range of possible fitness outcomes are diverse and span the parasitism-mutualism continuum. Contrary to previous predictions we found no evidence that transmission strategy, or the direct effects and indirect (plant-mediated) effects, of phytopathogens have divergent fitness outcomes for the vector. We discuss these findings in the context of plant – pathogen – vector ecology.

## Introduction

Many viral and bacterial pathogens that cause plant disease epidemics rely on herbivorous insect vectors for transmission (1,2). Vector-borne pathogens should be selected to enhance their transmission to plant hosts, via direct effects in the vector host or indirectly by manipulating the host plant (3). However, high virulence to the vector can negatively impact transmission as phytopathogens rely on the mobility of the vector for transmission and dispersal to non-motile plant hosts (4–6). Consequently, evolutionary theory predicts that vector-borne agents will be relatively less virulent to the vector, or even have beneficial phenotypes in the vector host (7–10). Despite predictions, a wide range of fitness effects have been reported across vector species. Squash vein yellowing virus (SqVYV) improves the longevity and fecundity of its whitefly vector (11), whereas Watermelon bud necrosis virus (WBNV) greatly reduces these fitness associated parameters in its vector *Thrips palmi* (12). Contradictory results have also been reported within the same taxa: *Candidatus* Liberibacter asiaticus positively affects the citrus psyllid’s fecundity (13,14), but negatively affects survival and longevity (14,15), underscoring the complexity of vector-pathogen interactions. The extent to which phytopathogens affect the fitness of vector hosts remains unquantified, despite the importance for the ecology and evolution of these pathosystems (a plant – pathogen – vector association) (3).

The outcome of vector-pathogen interactions may be affected by factors such as pathogen and vector taxonomy, differences in study methods, and whether vertical transmission occurs in the vector. However, most attention has focused on pathogen transmission mode, which describes the extent to which the pathogen replicates within the vector. Non-circulative pathogens are restricted to the vector’s stylet or foregut for short periods. Circulative viruses, and all bacteria, enter the haemocoel and render the vector infectious for longer periods (16), with some also propagating within the vector (17). Consequently, it has been suggested that non-circulative, non-persistent pathogens will predominantly have indirect effects on vector fitness, for example by affecting the volatile profile of infected plants (18) or altering plant defence against herbivory (19). In contrast, circulative, propagative viruses and bacteria are more likely to affect vector fitness directly, for example by hijacking the vector’s cellular machinery, in addition to plant-mediated indirect effects.

Previous vote counting studies suggest a positive effect of phytopathogens on their vectors (3,20,21); however, these focused either exclusively on viruses or included effects on vector behaviour. Here, we use a meta-analytic approach to test the hypothesis that vector-borne bacterial and viral phytopathogens benefit their insect vectors. We include only fitness-associated metrics to assess the scope for mutualism in these interactions. Further analysis tests for ecological drivers of variation in vector-pathogen interactions. Our synthesis of 34 plant – pathogen – vector systems suggests the mean effect of infection on vector fitness is neutral, but the range of outcomes is considerable and spans the parasitism-mutualism continuum. As fitness costs affect the abundance and population dynamics of insect vectors, an improved understanding of vector-pathogen interactions will be critical for preventing plant disease epidemics.

## Methods

### Literature Review

We searched Web of Science for relevant studies and retrieved 888 papers published up to July 2021, which were then screened before reading the full text and extracting data (Supplementary methods; supplementary figure 3). Studies on plant pathosystems were included when each experiment had a treatment and control group with independent samples; an infection challenge, where ≥70% of individuals were infected; a standardised way to quantify fitness; and ≥5 observations. Studies that evaluated the effects of temperature, co-infections, or starvation, and studies testing the effect of infection on host preference and feeding behaviours, were excluded. Overall, 26 studies were included covering 34 unique pathosystems.

### Calculation of effect sizes

Of the 115 effect sizes extracted, 85 were in count form and 30 were dichotomous. To combine the two datasets, dichotomous data were first expressed as odds ratios and accompanying SEs as described in (22), then converted into standardised mean differences (SMD) (23). Where count data was provided, the standardised mean difference was calculated as the difference in mean outcome between the infected and control groups, divided by the standard deviation of the outcome among participants. Where mean outcomes were not reported, data were obtained from plots by extracting data points using webplotdigitiser (24). Missing standard errors and sample sizes were extrapolated using methods described in (25).

### Meta-analytical model

Most studies reported multiple effect sizes. To account for any dependencies within studies, multi-level models were fitted using restricted maximum-likelihood estimation (REML) using the *metafor* package (v3.0-2) (26) in R v4.0.5 (27,28). Nesting the effects reported within the study ID allowed for differentiation of the effect sizes due to sampling variation within and between studies (29). Pathogen species, host species, and vector species were also included as random effects, allowing multiple representations of the same species to be accounted for (30). Taxonomic subgroup analysis was performed by calculating mean effect sizes for each pathogen genus and vector genus. The contribution of ecological and methodological predictors to the overall effect was then assessed using univariate models (Supplementary table 1). Omnibus tests were used to assess differences in mean effect size between groups, and likelihood ratio tests (LRT) using maximum likelihood estimation were used to assess the significance of each predictor.

### Assessing bias

Publication bias was visually assessed using funnel plots (Supplementary figure 4). The weighted Rosenberg method (31) was used to calculate a fail-safe number, which estimates the number of studies averaging a null result that would have to be added to the dataset to reduce the significance level to a target alpha level (e.g. 0.05). The fail-safe number was 581, which is not greater than the threshold considered for a robust analysis (nfs>5N + 10 where N = no. effect sizes). These results suggest that the dataset could be affected by bias towards positive results. However, as noted by others (20), negative results would be biologically interesting in this case, and less likely to remain unpublished.

## Results

### 1. Neutral effect of infection on vector host

Across diverse vector hosts and plant pathogen taxa, the mean effect of infection on the vector host was neutral, but highly variable (full model: SMD = -0.1354, SE = 0.3436, 95% CI: -0.8088, 0.5381, p = 0.6936) with interactions spanning from parasitic to mutualistic.

**Figure 1.**
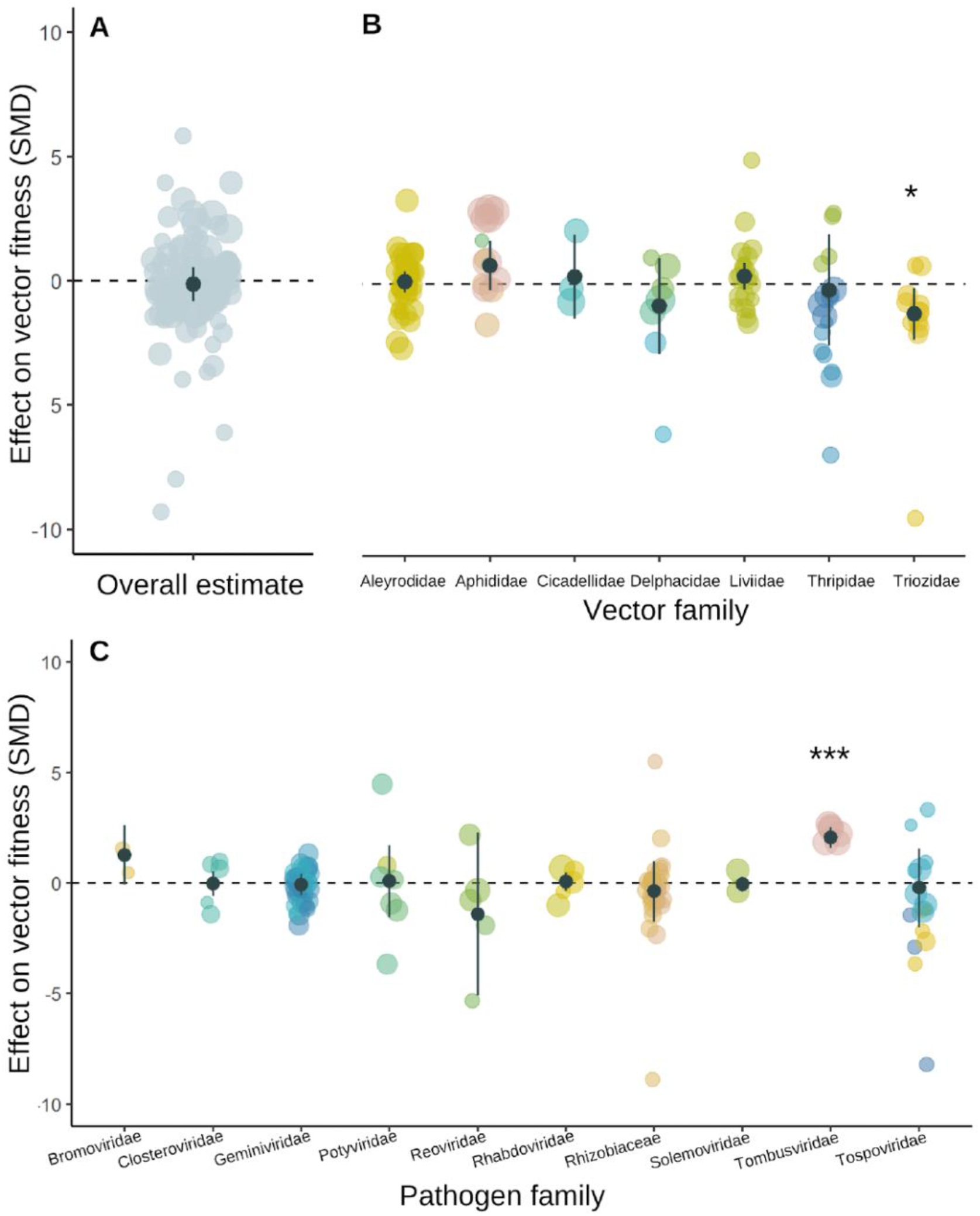
Vector-pathogen interactions are diverse and range from parasitic to mutualistic effects on vector fitness (SMD = -0.1354, 95% CI: -0.8088 -0.5381) (1A). Observational subgroup analysis according to vector family (1B) and pathogen family (1C) Asterisks denote significance (*p<0.5, ***p<0.001). Individual effect sizes are jittered and coloured by vector species, note the limited observations for some families. Point size represents study sample size. Mean effects and 95% confidence intervals are plotted in black.

**Figure 2.**
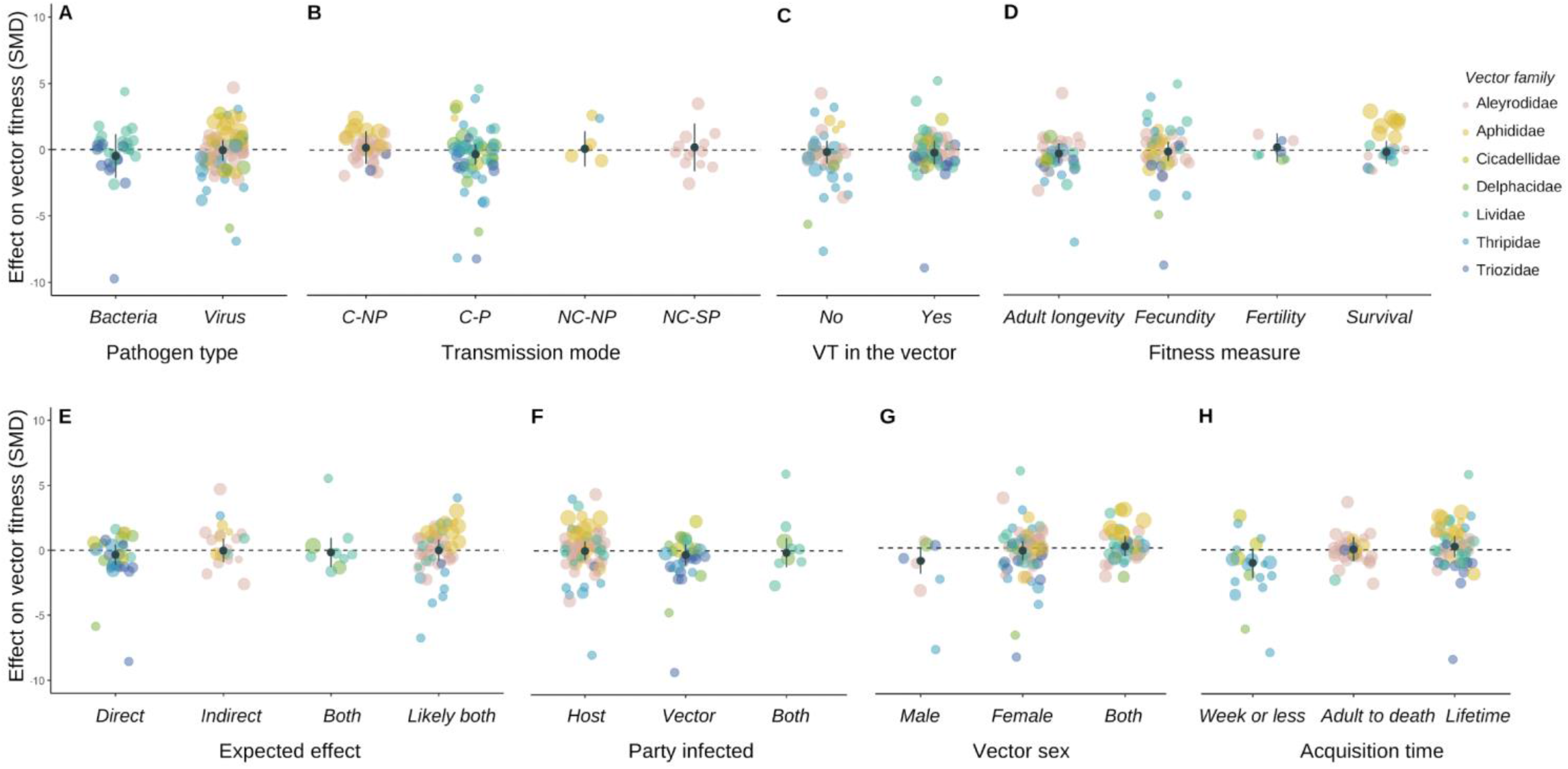
Variation in the effect of infection on vector fitness was not significantly affected by pathogen type (A), transmission mode (circulative non-propagative; circulative-propagative; non-circulative, non-persistent; non-circulative, semi-persistent) (B), vertical transmission in vector (C), fitness measure (D), direct or indirect effects (E), vector sex (F), party infected (G), or acquisition time (H). Individual effect sizes are jittered and coloured by vector family. Point size represents study sample size. Mean effect sizes and 95% confidence intervals are plotted in black.

**Table 1:**
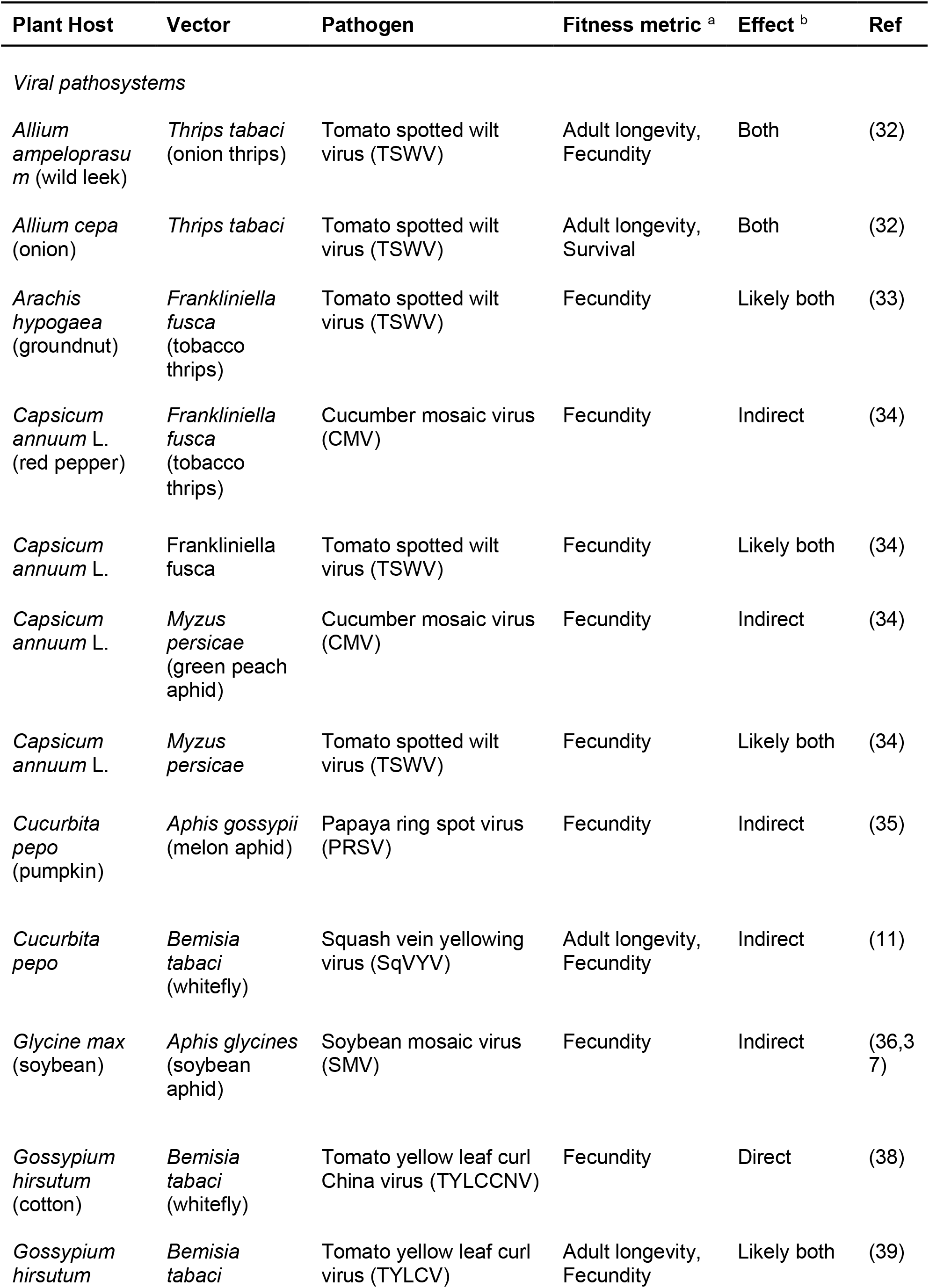
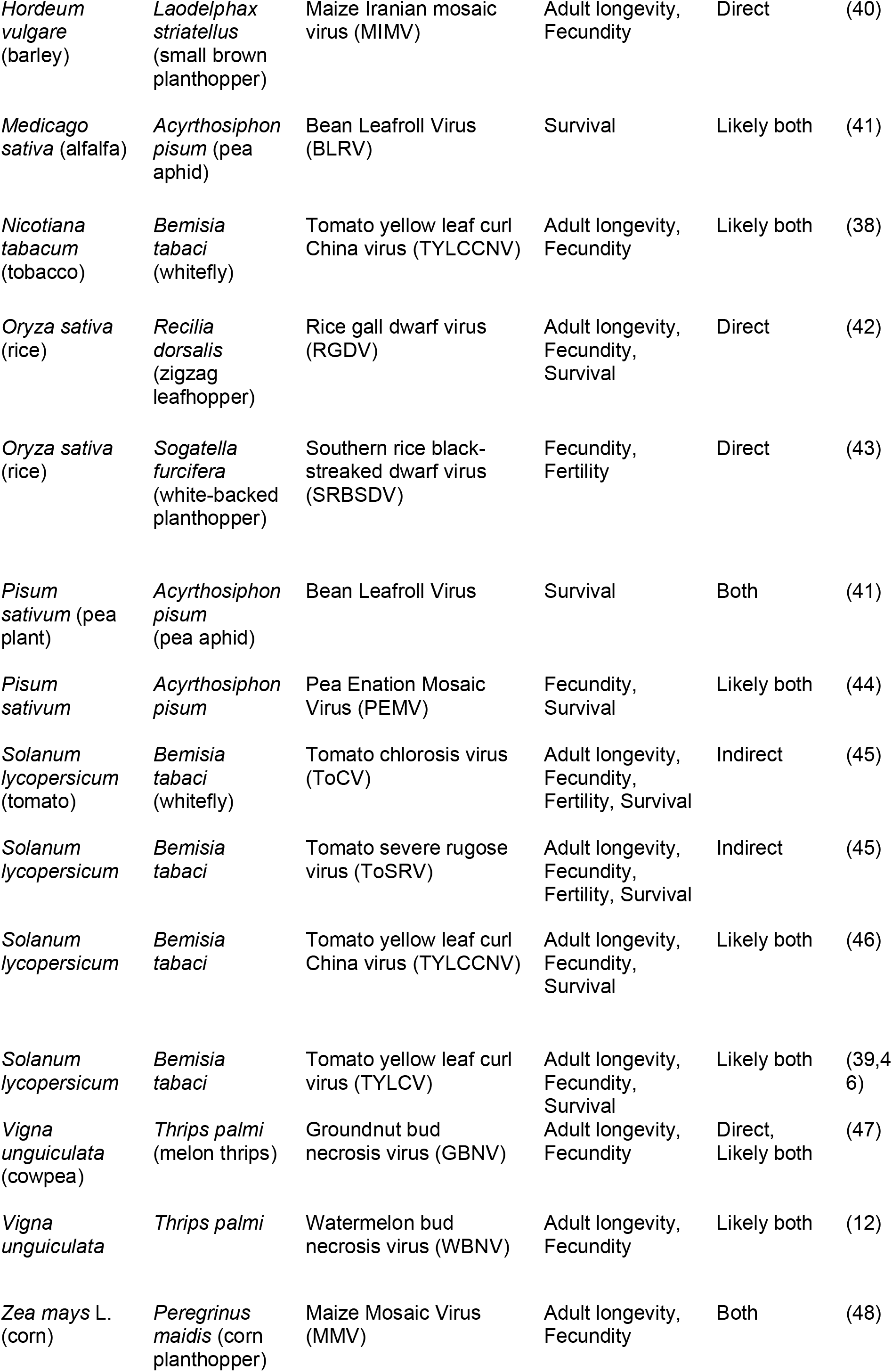
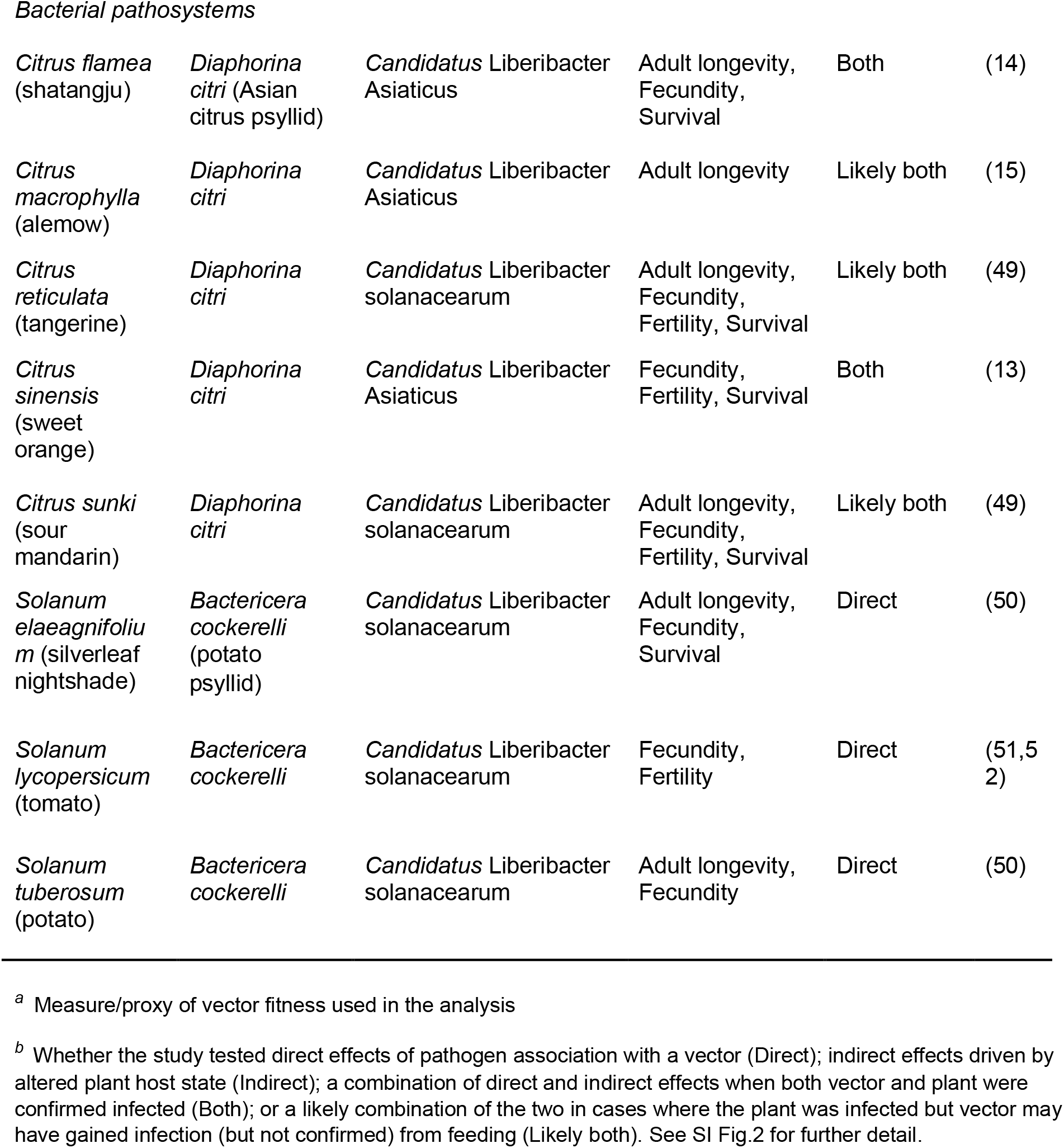
Summary of pathosystems included

| Plant Host | Vector | Pathogen | Fitness metric <sup>a</sup> | Effect <sup>b</sup> | Ref |
| --- | --- | --- | --- | --- | --- |
| *Viral pathosystems* |  |  |  |  |  |
| *Allium ampeloprasum* (wild leek) | *Thrips tabaci* (onion thrips) | Tomato spotted wilt virus (TSWV) | Adult longevity, Fecundity | Both | (32) |
| *Allium cepa* (onion) | *Thrips tabaci* | Tomato spotted wilt virus (TSWV) | Adult longevity, Survival | Both | (32) |
| *Arachis hypogaea* (groundnut) | *Frankliniella fusca* (tobacco thrips) | Tomato spotted wilt virus (TSWV) | Fecundity | Likely both | (33) |
| *Capsicum annuum* L. (red pepper) | *Frankliniella fusca* (tobacco thrips) | Cucumber mosaic virus (CMV) | Fecundity | Indirect | (34) |
| *Capsicum annuum* L. | *Frankliniella fusca* | Tomato spotted wilt virus (TSWV) | Fecundity | Likely both | (34) |
| *Capsicum annuum* L. | *Myzus persicae* (green peach aphid) | Cucumber mosaic virus (CMV) | Fecundity | Indirect | (34) |
| *Capsicum annuum* L. | *Myzus persicae* | Tomato spotted wilt virus (TSWV) | Fecundity | Likely both | (34) |
| *Cucurbita pepo* (pumpkin) | *Aphis gossypii* (melon aphid) | Papaya ring spot virus (PRSV) | Fecundity | Indirect | (35) |
| *Cucurbita pepo* | *Bemisia tabaci* (whitefly) | Squash vein yellowing virus (SqVYV) | Adult longevity, Fecundity | Indirect | (11) |
| *Glycine max* (soybean) | *Aphis glycines* (soybean aphid) | Soybean mosaic virus (SMV) | Fecundity | Indirect | (36,37) |
| *Gossypium hirsutum* (cotton) | *Bemisia tabaci* (whitefly) | Tomato yellow leaf curl China virus (TYLCCNV) | Fecundity | Direct | (38) |
| *Gossypium hirsutum* | *Bemisia tabaci* | Tomato yellow leaf curl virus (TYLCV) | Adult longevity, Fecundity | Likely both | (39) |
| *Hordeum vulgare* (barley) | *Laodelphax striatellus* (small brown planthopper) | Maize Iranian mosaic virus (MIMV) | Adult longevity, Fecundity | Direct | (40) |
| *Medicago sativa* (alfalfa) | *Acyrtosiphon pisum* (pea aphid) | Bean Leafroll Virus (BLRV) | Survival | Likely both | (41) |
| *Nicotiana tabacum* (tobacco) | *Bemisia tabaci* (whitefly) | Tomato yellow leaf curl China virus (TYLCCNV) | Adult longevity, Fecundity | Likely both | (38) |
| *Oryza sativa* (rice) | *Recilia dorsalis* (zigzag leafhopper) | Rice gall dwarf virus (RGDV) | Adult longevity, Fecundity, Survival | Direct | (42) |
| *Oryza sativa* (rice) | *Sogatella furcifera* (white-backed planthopper) | Southern rice black-streaked dwarf virus (SRBSDV) | Fecundity, Fertility | Direct | (43) |
| *Pisum sativum* (pea plant) | *Acyrtosiphon pisum* (pea aphid) | Bean Leafroll Virus | Survival | Both | (41) |
| *Pisum sativum* | *Acyrtosiphon pisum* | Pea Enation Mosaic Virus (PEMV) | Fecundity, Survival | Likely both | (44) |
| *Solanum lycopersicum* (tomato) | *Bemisia tabaci* (whitefly) | Tomato chlorosis virus (ToCV) | Adult longevity, Fecundity, Fertility, Survival | Indirect | (45) |
| *Solanum lycopersicum* | *Bemisia tabaci* | Tomato severe rugose virus (ToSRV) | Adult longevity, Fecundity, Fertility, Survival | Indirect | (45) |
| *Solanum lycopersicum* | *Bemisia tabaci* | Tomato yellow leaf curl China virus (TYLCCNV) | Adult longevity, Fecundity, Survival | Likely both | (46) |
| *Solanum lycopersicum* | *Bemisia tabaci* | Tomato yellow leaf curl virus (TYLCV) | Adult longevity, Fecundity, Survival | Likely both | (39,46) |
| *Vigna unguiculata* (cowpea) | *Thrips palmi* (melon thrips) | Groundnut bud necrosis virus (GBNV) | Adult longevity, Fecundity | Direct, Likely both | (47) |
| *Vigna unguiculata* | *Thrips palmi* | Watermelon bud necrosis virus (WBNV) | Adult longevity, Fecundity | Likely both | (12) |
| *Zea mays* L. (corn) | *Peregrinus maidis* (corn planthopper) | Maize Mosaic Virus (MMV) | Adult longevity, Fecundity | Both | (48) |
| Bacterial pathosystems |  |  |  |  |  |
| *Citrus flamea*<br>(shatangju) | *Diaphorina citri* (Asian citrus psyllid) | *Candidatus Liberibacter Asiaticus* | Adult longevity, Fecundity, Survival | Both | (14) |
| *Citrus macrophylla*<br>(alemow) | *Diaphorina citri* | *Candidatus Liberibacter Asiaticus* | Adult longevity | Likely both | (15) |
| *Citrus reticulata*<br>(tangerine) | *Diaphorina citri* | *Candidatus Liberibacter solanacearum* | Adult longevity, Fecundity, Fertility, Survival | Likely both | (49) |
| *Citrus sinensis*<br>(sweet orange) | *Diaphorina citri* | *Candidatus Liberibacter Asiaticus* | Fecundity, Fertility, Survival | Both | (13) |
| *Citrus sunki*<br>(sour mandarin) | *Diaphorina citri* | *Candidatus Liberibacter solanacearum* | Adult longevity, Fecundity, Fertility, Survival | Likely both | (49) |
| *Solanum elaeagnifolium*<br>(silverleaf nightshade) | *Bactericera cockerelli* (potato psyllid) | *Candidatus Liberibacter solanacearum* | Adult longevity, Fecundity, Survival | Direct | (50) |
| *Solanum lycopersicum*<br>(tomato) | *Bactericera cockerelli* | *Candidatus Liberibacter solanacearum* | Fecundity, Fertility | Direct | (51,52) |
| *Solanum tuberosum*<br>(potato) | *Bactericera cockerelli* | *Candidatus Liberibacter solanacearum* | Adult longevity, Fecundity | Direct | (50) |

<sup>a</sup> Measure/proxy of vector fitness used in the analysis
<sup>b</sup> Whether the study tested direct effects of pathogen association with a vector (Direct); indirect effects driven by altered plant host state (Indirect); a combination of direct and indirect effects when both vector and plant were confirmed infected (Both); or a likely combination of the two in cases where the plant was infected but vector may have gained infection (but not confirmed) from feeding (Likely both). See SI Fig.2 for further detail.

Heterogeneity across studies was very high (Q(df = 114) = 1090.6284, p-val < .0001), a common finding in ecological and evolutionary meta-analyses (53).

### 2. Subgroup analysis according to phylogeny

Mean effects were calculated for taxonomic subgroups at the family level and most showed neutral effects overall. However, infections in the vector family Triozidae (jumping plant lice) were associated with a negative impact on vector host fitness, (SMD= -1.2066, SE = 0.5329, 95% CI = -2.2511, -0.1621, p= 0.0236). Conversely, infections in the pathogen family Tombusviridae (single-stranded RNA viruses) conferred a positive effect on vector fitness (SMD= 2.0543, SE= 0.2404, 95% CI= 1.5831, 2.5255, p= <.0001).

### 3. Biological predictors

We tested multiple ecological factors to assess if they accounted for variation in the effect of infection on vector fitness. Fitness effects did not vary between bacterial and viral pathogens (QM(df = 1) = 0.2041, p-val = 0.6514), and transmission mode did not significantly affect the fitness of the vector host (QM(df = 3) = 0.7933, p-val = 0.8511). Estimates for vector fitness were not impacted by pathogens that are also transmitted vertically in the vector (horizontal/mixed-mode transmission, QM(df = 1) = 0.0090, p-val = 0.9244). We did not detect significant differences between the measures used to estimate fitness (e.g., longevity, fecundity, fertility, and survival (QM(df = 3) = 1.0689, p-val = 0.7846).

Pathosystems were classified by whether the infections were expected to directly affect fitness as they infect the vector host (Direct effects); indirectly affect fitness by altering plant host defences to herbivory and/or facilitating predation (Indirect effects); a certain combination of direct and indirect effects (Both); and a likely combination of the two (Likely both). For full classification criteria see supplementary figure 2. We were unable to detect significant differences in the fitness outcomes conferred by infections across these categories (QM(df = 3) = 0.6531, p-val = 0.8842).

Overall, sex did not have a significant effect on the fitness of infected vectors (QM(df = 2) = 5.5433, p-val = 0.0626). However, infected males were generally more negatively affected by infection (SMD = -0.9706, 95% CI = -1.9570, 0.0159) than females (SMD = -0.2159, 95% CI = -0.9220, 0.4902) or mixed groups (SMD = 0.1208, 95% CI = -0.6425, 0.8842).

### 4. Methodological predictors

We found limited evidence that methodological factors influenced the effect of infection on vector host fitness. The method of pathogen inoculation to the vector host did not significantly affect fitness outcomes (QM(df = 2) = 0.2068, p-val = 0.9018), and for studies that inoculated the plant host no effect of inoculation method was detected (QM(df = 3) = 2.2931, p-val = 0.5138).The parties experimentally infected during the study also did not have a significant impact on the infection outcome (QM(df = 2) = 0.6016, p-val = 0.7402). However, vector fitness was more negatively affected in studies where both plant and vector hosts, or only the vector host was infected (SMD = -0.1619, 95% CI = -1.276, 0.9522; SMD = -0.3198, 95% CI = -1.1064, 0.4668), whereas vectors showed neutral fitness effects in studies where only the plant host was infected (SMD = -0.0189, 95% CI = -0.7388, 0.7011).

Finally, the amount of time allotted for the vector to acquire the pathogen (acquisition time) did not significantly affect the outcome of infection (QM(df = 2) = 3.7372, p-val = 0.1543). That said, vectors who had shorter acquisition periods of a week or less generally had reduced fitness (SMD = -1.0049, 95% CI = -2.1278, 0.118) but those with longer acquisition periods (adult to death/lifetime) showed neutral fitness effects (SMD = 0.0542, 95% CI = -0.8832, 0.9917; SMD = 0.2564, 95% CI = -0.5433, 1.0561).

## Discussion

Evolutionary theory suggests that vector-borne pathogens will be selected for low virulence or even beneficial phenotypes in the vector host, as mobility of this host is crucial for transmission (5,6,54). Across 34 pathosystems we show the average effect of infection on vector fitness is neutral, in line with theoretical models (7–10). However, our analysis shows vector-pathogen interactions span the symbiosis continuum, with infections ranging from beneficial to highly detrimental for vectors.

Phylogeny is frequently an important predictor of host-microbe interactions (55,56). Our analysis revealed that among pathogen families, the Tombusviridae (single stranded plant RNA viruses) significantly increase vector fitness. Among vector families (Supplementary figure 1), the jumping lice Triozidae were notable in being more negatively affected by infection. These results provisionally suggest that phylogeny, or shared ecology, may play a role in infection outcomes; however, the mechanisms through which phytopathogens harm or benefit their vectors are not well-understood (3). Sex-specific differences can also drive variation in host-parasite interactions (57), and in our study male vectors appeared to suffer more from infections. Here, the negative trend for males may be influenced by differences in immunocompetence (58); alternatively, less harm may be favoured in females who are generally the dominant transmitters (59).

Contrary to our initial hypothesis, circulative pathogens had similar fitness effects on vectors to those that do not circulate within the host. Similarly, we found no evidence that direct effects of infection differ significantly from indirect effects. These results come with a caveat: vectors typically acquire pathogens by feeding on infected plants, making it challenging to disentangle the direct and indirect effects of the pathogen on the insect vector. Some studies limited the contribution of indirect effects (14,38,40,42,43,47,50–52) by infecting vectors in vitro or transferring them regularly onto unexposed plants (47). For most other studies, whether effects were direct, indirect, likely both, or certainly both was inferred based on pathogen transmission mode and the experimentally infected party (supplementary figure 2). More studies are urgently needed that can distinguish the direct and indirect effects of plant pathogens on their vectors and establish whether mutualistic phenotypes evolve more readily within one of these effect classes.

Previously, some of the large variation in vector fitness effects has been attributed to differences in methods (60). For example, plant hosts inoculated mechanically vs vector inoculated will not induce processes such as herbivore effector triggered immunity or other changes to plant chemical composition (61–63).However, inoculation method did not explain much of the variation in vector fitness, suggesting these methods are more comparable than previously assumed. Notably, negative fitness outcomes were slightly more prevalent if the vector was the only party infected in the experiment. This suggests that beneficial phenotypes in the vector may depend upon the interaction of both direct and plant-mediated (indirect) effects of a pathogen. Disentangling the contribution of direct and indirect effects is difficult and will require more studies that infect each party individually and in combination.

Our work formally quantifies the effect of diverse phytopathogens on insect vectors. We report an overall neutral fitness effect, but one that is underpinned by considerable diversity in both parasitic and mutualistic phenotypes. This finding highlights the value of vector-phytopathogen systems for exploring the evolution of symbiotic diversity in three-player communities and highlights the necessity for pathosystem-specific approaches to vector control.

## Acknowledgements

This work was funded by a European Starter Grant (COEVOPRO 802242) to K.C.K. and a Jardine Scholarship to M.F.S. We thank Kim Hoang and Jingdi Li for sharing code, and the authors who provided access to their data. The authors declare no competing interests.

## Data accessibility

All data and code associated with this draft manuscript is publicly available in the fig share repository - https://doi.org/10.6084/m9.figshare.c.6215267.v1

